# A benchmarking framework for perturbational signature recontextualization across biological contexts

**DOI:** 10.64898/2026.08.14.744937

**Authors:** Andrew Chen, Noa Rappaport, Nick Schork, Thomas Girke, Stefano Monti

## Abstract

Perturbational transcriptomics is a powerful tool for understanding gene function and drug effects, yet predicting how perturbations manifest across different biological contexts remains a central challenge, limiting translation from model systems to clinically relevant tissues. Despite growing interest in this problem, benchmarking efforts have been hindered by inconsistent evaluation tasks, heterogeneous metrics, and limited assessment across perturbation types and biological systems. Here, we introduce a benchmarking framework for cross-context perturbation-signature prediction (a task we define as *signature recontextualization*), grounded in explicit definitions of the prediction task, target-data availability, and evaluation metrics centered on signature recovery. The framework evaluates prediction performance across three target-context data regimes: (1) control only, where only control profiles from the target context are measured; (2) low coverage, where a limited subset of perturbations in the target context are measured; and (3) high coverage, where most perturbations in the target context are measured. This design enables systematic assessment of how prediction performance depends on target-context sample size while providing a standardized basis for comparing methods. We evaluate newly developed projection-based (projectCor) and network-based (netProp) methods alongside deep learning-based foundation models (*sc*GPT, STACK) and statistical baselines. The benchmark spans diverse perturbational datasets: CRISPR knockdowns and drug perturbations in cell lines, plus in vivo chemical perturbations in rat tissues from DrugMatrix, extending evaluation beyond isolated cell-line models to tissue-level responses. For each method, evaluation was limited to the datasets, and perturbation types each model supports. Across tasks, projection and network propagation approaches show strong flexibility across perturbation types and biological contexts, and in several cases match or exceed the performance of deep learning and foundation models. We further show that perturbation predictability varies substantially with pathway conservation, transcriptional response strength, and baseline similarity between source and target contexts. All datasets, methods, and evaluation utilities are released as an open-source R package (*sigRecon*), providing a foundation for reproducible benchmarking and future method development.

## Introduction

Our ability to conduct large-scale, multiplexed perturbational screens has greatly increased in the past decade. Technologies such as genome-wide perturb-seq and combinatorial molecular barcoding, combined with advances in single-cell RNA sequencing, have enabled the systematic profiling of transcriptional responses to thousands of genetic and pharmacologic interventions across diverse cell types and tissues. Landmark resources such as the Connectivity Map (CMap) ^1^, Tahoe 100M ^2^, X-atlas ^3^, and other genome-wide CRISPR screens ^4–6^ have collectively produced datasets of unprecedented scale, generating rich catalogs of how gene expression is perturbed in response to both genetic and chemical stimuli. These growing repositories of perturbational omics data have served as foundational resources in functional genomics, greatly accelerating data-driven approaches to understanding gene function, drug mechanism of action, and cellular state transitions.

Motivated by this abundance of perturbational data, a large body of work has also emerged aiming to build predictive models of transcriptional responses to unseen perturbations. Early approaches based on linear models and gene regulatory networks have since been complemented by deep learning architectures capable of learning complex nonlinear relationships between perturbations and gene expression profiles. More recently, the advent of large-scale foundation models pretrained on massive corpora of transcriptomic data such as scGPT ^7^, Geneformer ^8^, scFoundation ^9^, CellFM ^10^, STATE ^11^, and STACK ^12^, or hybrid methods such as txPert which combines deep learning architectures with biological knowledge graphs ^13^, have promised to push the boundaries of perturbation response prediction further, with the prospect that such models could serve as core components of an in silico “virtual cell”^14^.

Despite this enthusiasm, recent empirical evaluations have cast doubt on whether current deep learning and foundation models live up to this promise. Several independent benchmarking studies ^15–23^ as well as community-organized competitions ^24,25^ have revealed a sobering pattern: deep learning and foundation models have yet to consistently outperform simple baselines for perturbation response prediction, such as predicting the mean profile of all training perturbations as the prediction. These findings suggest that, while the field has made significant progress, critical gaps remain in our ability to generalize perturbation predictions reliably. These gaps are not merely a matter of model capacity; they reflect deeper challenges in how models are trained, evaluated, and deployed across the diverse range of experimental contexts encountered in practice. Given the results of these prior benchmarking studies, we constrained the choice of foundation models to be included in our benchmarking to scGPT and STACK. Other methods such as STATE were omitted since its pretraining included the same perturb-seq and Tahoe perturbational datasets that are used in benchmarking.

A currently underexplored aspect of perturbation response prediction is its connection to a specific yet practically relevant experimental scenario: a researcher knows the transcriptional signature of a perturbation in one biological context – for example, in a cell line or in vitro system – and wishes to predict what that same perturbation would look like in another context, such as a different cell line, a primary tissue, or an in vivo setting. We refer to this task throughout this study as *signature recontextualization*. While other studies have benchmarked methods predicting transcriptomic responses to perturbations across covariates such as cell type, timepoints, and donors ^16,23,26–29^, to our knowledge no existing benchmarking framework has treated recovery of the differentially expressed gene (DEG) signature as the evaluation target across both genetic and chemical perturbations while varying target-context data availability, despite its clear relevance to translational research and the design of experimental follow-up studies.

Here, we introduce a rigorous benchmarking framework centered on recontextualizing perturbational signatures across biological contexts and make several key contributions to the field (Fig. 1). First, we systematically evaluate prediction performance as a function of target-context sample size – a dimension largely unexplored in prior benchmarking efforts. We do this by defining three benchmarking regimes that span the spectrum of target-context data availability: (1) a control regime, where only control expression profiles from the target context are available; (2) a low coverage regime, where a limited set (1/10^th^) of perturbations has been measured in the target context; and (3) a high coverage regime, where a substantial set (9/10^th^) of perturbations in the target context is available as prior knowledge. By evaluating across these regimes, we construct a partial learning curve for each method and dataset pair, revealing how prediction accuracy scales with target-context training data. While existing benchmarking efforts have focused almost exclusively on the high coverage regime – implicitly assuming abundant target-context data – the control and low coverage regimes reflect situations routinely encountered in practice, particularly when exploring new biological contexts where experimental resources are limited (Box 1).

**Figure 1.**
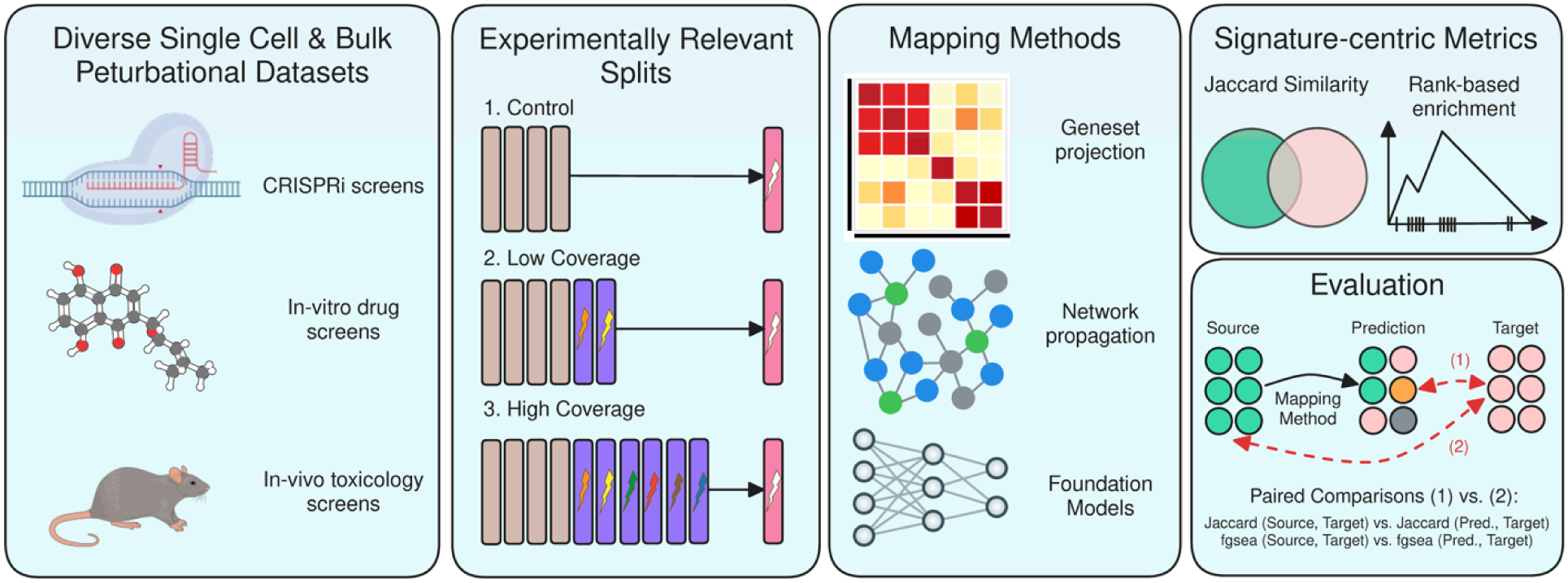
Overview of datasets, benchmarking strategies, methods, and evaluation metrics

#### Box 1 Benchmarking Regimes

We evaluate prediction performance across three sample-size regimes, effectively measuring a three-point learning curve for each method-dataset pair. Each regime is motivated by realistic laboratory scenarios:

##### 1. Control Regime

*<u>Setup:</u>* Given perturbed profiles in context Y, and *<u>only</u>* unperturbed profiles in context Z, we predict gene signatures of the same perturbations in context Z.

*<u>Scenario</u>:* A cancer researcher has conducted and derived perturbational signatures of drug perturbations in ER+ MCF7 cell line but wants to estimate the same signatures in a triple-negative MDA-MB-231 cancer cell line given only unperturbed profiles of MDA-MB-231 from the cancer cell line encyclopedia. This reflects pure cross-context transfer with no target-context labeled data.

##### 2. Low Coverage

*<u>Setup:</u>* Given perturbed profiles in context Y, and 10% *<u>of the measured perturbations</u>* in context Z, we predict gene signatures of held-out unmeasured perturbations in context Z.

*<u>Scenario</u>*: A CRISPR researcher has performed genome-wide perturb-seq experiments in CD4 positive T-cells but only has enough resources to perform 10% of the same gene perturbations in CD8 positive T-cells. The sparse perturbational data in CD8 positive T-cells is used to train a model to predict perturb-seq signatures for the unmeasured genes.

##### 3. High Coverage

*<u>Setup:</u>* Given perturbed profiles in context Y, and 90% *<u>of the measured perturbations</u>* in context Z, we predict gene signatures of held-out unmeasured perturbations in context Z.

*<u>Scenario</u>*: A computational researcher is training a foundation model on all publicly available single cell drug perturbational data collected on GEO for the purposes of predicting unseen perturbations in new biological contexts. This reflects the implicit assumption of prior benchmarking work: that extensive target-context data is available.

Second, we substantially expand the scope of perturbation response prediction beyond cell line settings. Our benchmark includes cross-tissue bulk transcriptomic drug responses from *DrugMatrix* ^30^, extending prior efforts beyond isolated genetic and chemical perturbations in cell lines to encompass in vivo drug responses, a more translationally representative setting.

Third, alongside these deep learning and foundation model comparisons, we develop and evaluate two new methods: projectCor, a signature projection approach that maps perturbational profiles from a source context onto a target context via expression correlation structure, and a network propagation method (netProp) that leverages gene network topology to predict new perturbational signals across biological contexts. These methods are computationally lightweight and, unlike existing foundation models, broadly applicable across perturbation types and input data modalities. We show that these approaches were competitive with state-of-the-art deep learning and foundation models, outranking them in *Perturb-Seq and SciPlex* (*sc*GPT was evaluated only on Perturb-Seq and STACK on three of four datasets), consistent with recent evidence that model complexity does not inherently improve generalization in perturbation prediction tasks.

Together, these contributions – systematic learning-curve analysis, expanded perturbation types and biological contexts, and the development of flexible baseline methods – provide a rigorous foundation for evaluating perturbation response prediction across biological contexts. To facilitate reproducibility and community adoption, we release all benchmark datasets, method implementations, and evaluation utilities as an open-source R package, providing a comprehensive and extensible framework for future method development in perturbational prediction.

## Results

### Unperturbed profiles are insufficient for existing methods

The control benchmarking regime evaluates whether methods can leverage unperturbed baseline expression profiles in the target context to predict perturbed expression profiles in the target context. Across all datasets, no method reliably performs better than the no-change baseline as seen through the median delta normalized enrichment score (NES) and delta Jaccard score (Jacc) being below the no-change line (Fig. 2).

**Figure 2.**
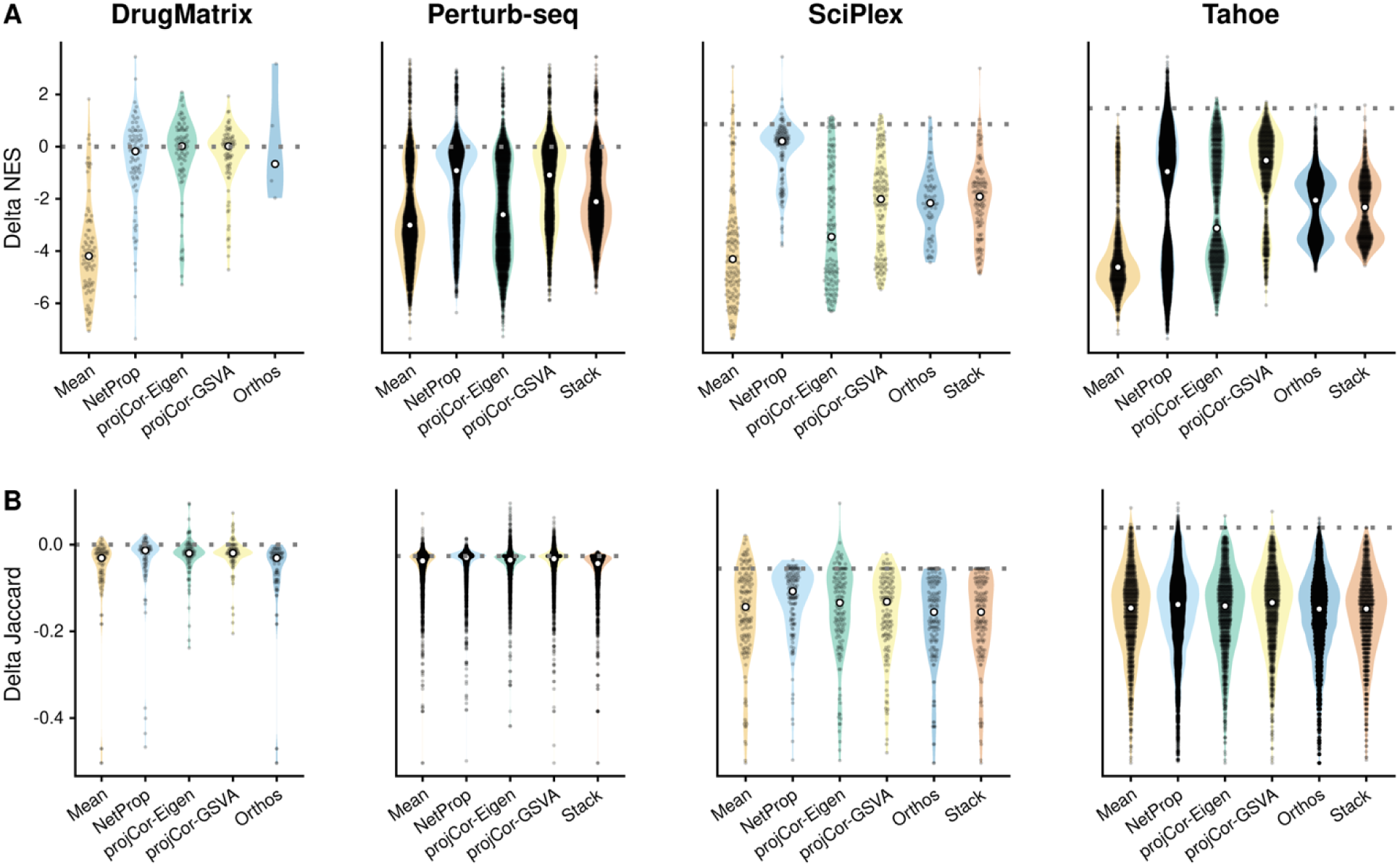
Performance of methods in the control regime. A) Difference in NES between the No-Change baseline and each other method. B) Difference in Jaccard similarity between the No-Change baseline and each other method.

While the overall conclusion from the control benchmarking results is consistent across methods, the exact mechanisms for which each method leverages unperturbed data varies: projectCor and netProp leverage input data to estimate covariance, which for unperturbed profiles, largely reflects homeostatic baseline variation rather than perturbation-specific regulatory structure, whereas Orthos and STACK leverage input data to decompose perturbation specific effects, and condition context-specific generation respectively.

### With perturbed profiles, projection and network-based methods perform favorably

Once perturbed profiles were incorporated, either a 10% split (low coverage regime) or a 90% split of target perturbations (high coverage regime), many prediction methods perform better than the no-change baseline (Fig. 3). For all results in the low coverage and high coverage regimes, we report median performance across all possible splits of perturbed data (Supp. Table 1a, Supp. Fig. 1,2)

**Figure 3.**
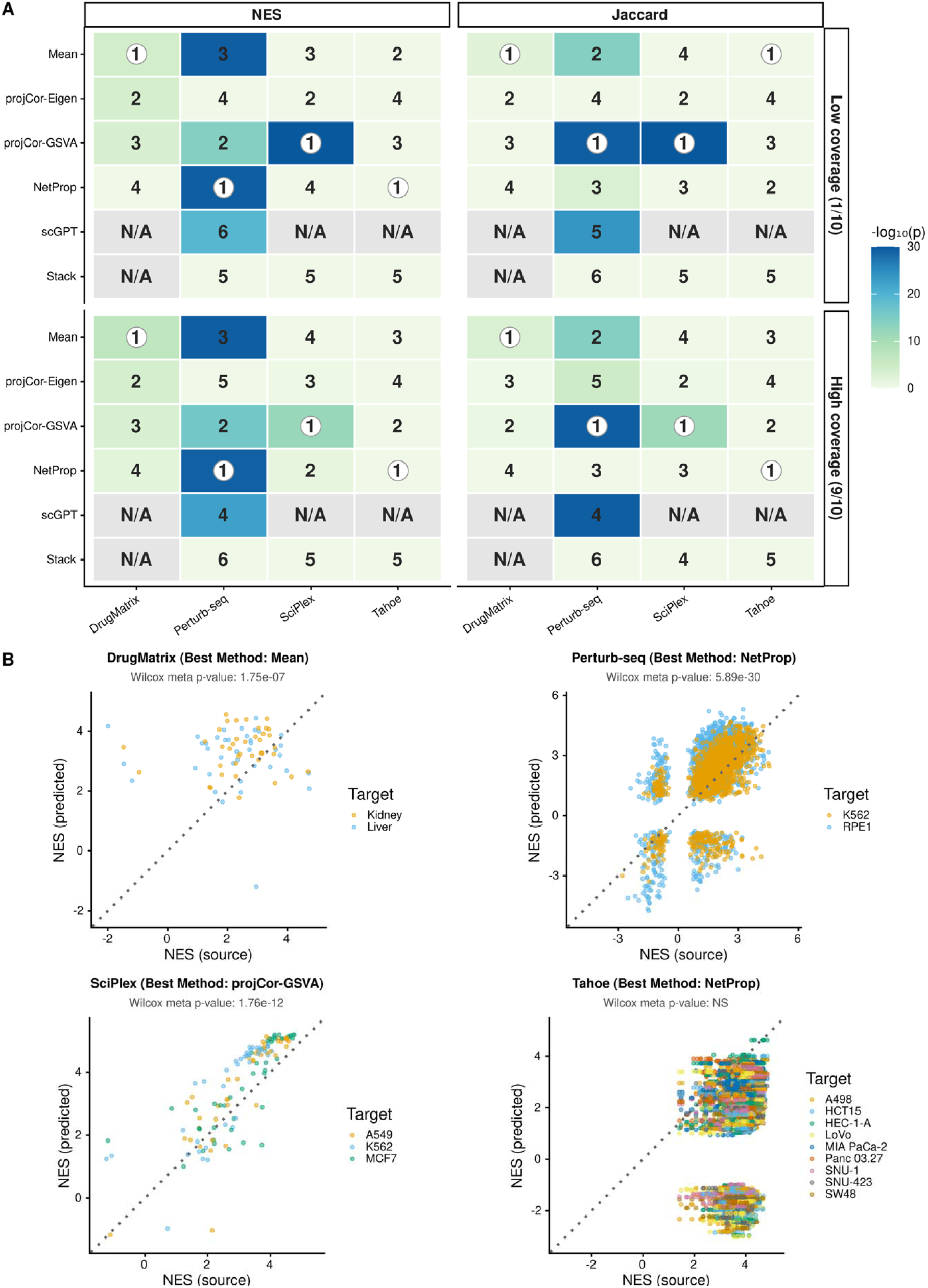
A) Top-ranked method according to a one-sided paired Wilcoxon test (Method X > No-Change) using both NES and Jaccard similarity. Cell shading is the fisher’s p.value, and the rank is computed according to the Wilcoxon test statistic. B) NES before and after recontextualization, for the top-ranked method in each dataset.

**Table 1:** Summary of Perturbational Datasets.

| Dataset | Year | System | # Sig. Perts. | # Contexts | Sig. Derivation |
| --- | --- | --- | --- | --- | --- |
| *DrugMatrix* | 2014 | Affymetrix GeneChip | 78 | 2 | Limma |
| *SciPlex* | 2019 | Illumina NovaSeq 6000 | 129 | 3 | Pseudobulk + DESeq2 |
| *Perturb-Seq* | 2022 | Illumina NovaSeq 6000 | 2557 | 2 | Pseudobulk + DESeq2 |
| *Tahoe* | 2025 | Ultima Genomics UG100 | 875 | 10 | Pseudobulk + DESeq2 |

For the *DrugMatrix* dataset, mean, projCor-GSVA, projCor-eigen and netProp all significantly improved upon the no-change baseline in terms of the NES metric, but only the mean baseline significantly improved signature similarity in terms of the Jaccard metric (Supp. Table 1bc). Overall, the mean baseline was the highest ranked method for recontextualizing *DrugMatrix* perturbations (Fig. 3a,b, Supp. Fig. 3).

For the *Perturb-Seq* dataset, mean, projCor-GSVA, netProp, and scGPT all significantly improved upon the no-change baseline in terms of NES and Jaccard similarity (Supp. Table 1bc). Overall, netProp is the highest ranked method for both benchmarking regimes in terms of NES, but projCor-GSVA is the best method for both perturbed regimes under the lens of Jaccard similarity (Fig. 3a,b, Supp. Fig. 3).

For the *SciPlex* dataset, projCor-GSVA and netProp significantly improved signatures post-recontextualization under the NES metric (Supp. Table 1bc), but only the former improved signatures relative to the no-change baseline given the Jaccard metric (Fig. 3a,b, Supp. Fig. 3).

For the *Tahoe* dataset, no method improved upon the no-change baseline in either regime, or with either metric (Fig. 3a,b, Supp. Fig. 3).

### Not all perturbations are equal

Since these perturbational datasets report transcriptomic changes for a diverse range of perturbations and biological contexts (up to 1000 perturbations across 50 cell-lines for *Tahoe*), we also investigated which types of perturbations are consistently harder or more easily mapped across contexts. To categorize types of perturbations, we grouped drug perturbations according to their targets/mechanism of action (MOA) (Supp. Table 2), and utilized statistical measures of perturbational impact from the *TRADE* package ^5^.

#### Mechanism of Action (MOA)

For each of the three drug perturbation datasets (*DrugMatrix, SciPlex, Tahoe-100M*), we observed that recontextualization performance varies across drug target/MOA class (Fig. 4, Supp. Fig. 6, Supp. Fig. 7). For DrugMatrix, Statins and fibrates, which act largely through shared SREBP2- and PPARα-regulated lipid-metabolism circuitry that is broadly conserved across liver and kidney, showed among the largest positive Delta NES values in our benchmark, consistent with convergent transcriptional regulation of these pathways across tissues ^31–33^. In contrast, corticosteroids, which act through a glucocorticoid receptor (GR) cistrome that is tissue-specific ^34^, and anthracyclines, whose activity depends on TOP2B-mediated DNA damage in a chromatin-context-dependent manner ^35,36^, tended to show poorer transfer, consistent with mechanisms that depend on tissue-specific regulatory or chromatin context rather than on a pathway that is broadly conserved across tissues.

**Figure 4.**
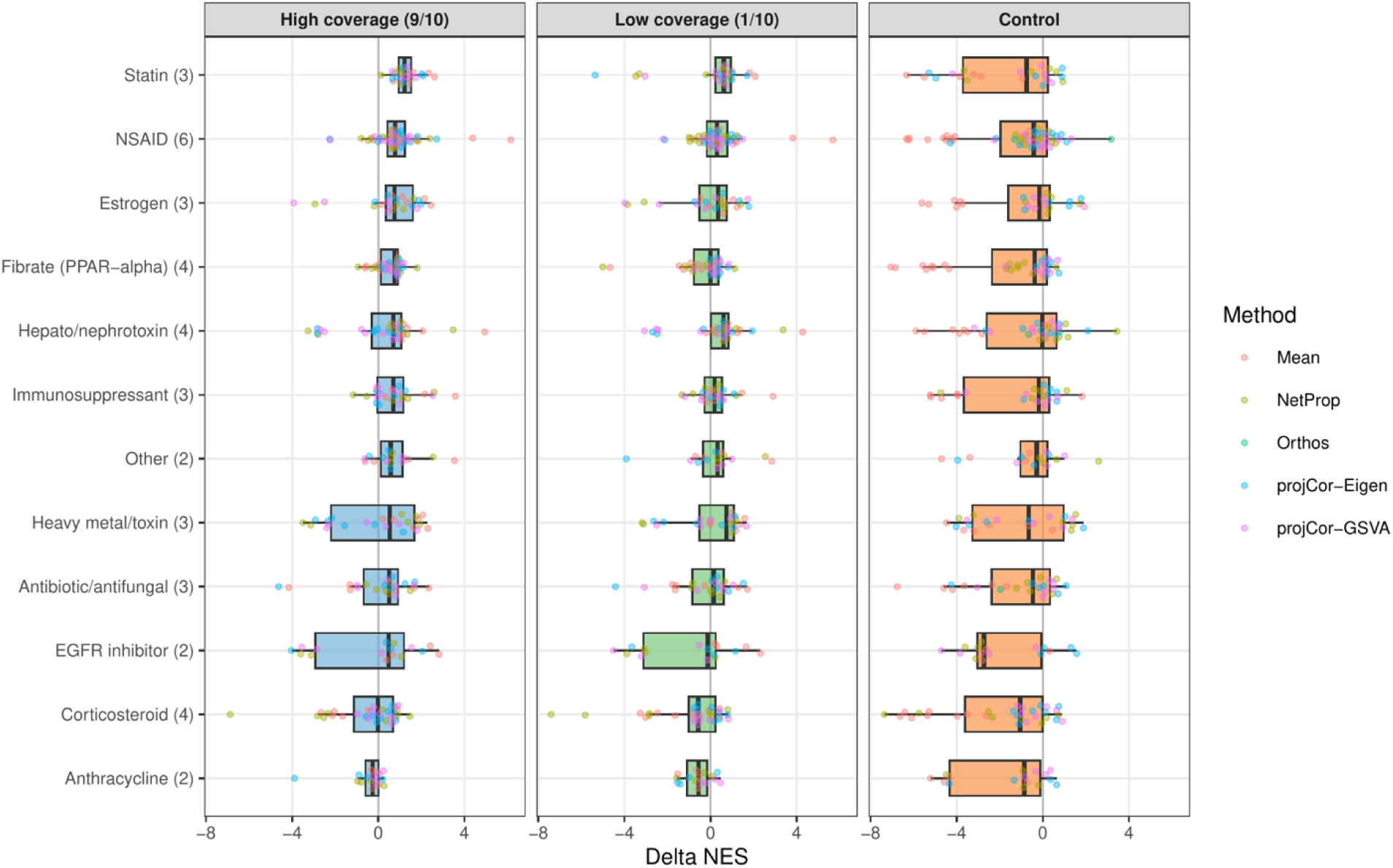
Delta NES (prediction - no change), stratified by drug mechanism of action as well as benchmarking regime, across all methods and target tissues (kidney, liver) in DrugMatrix.

**Figure 5.**
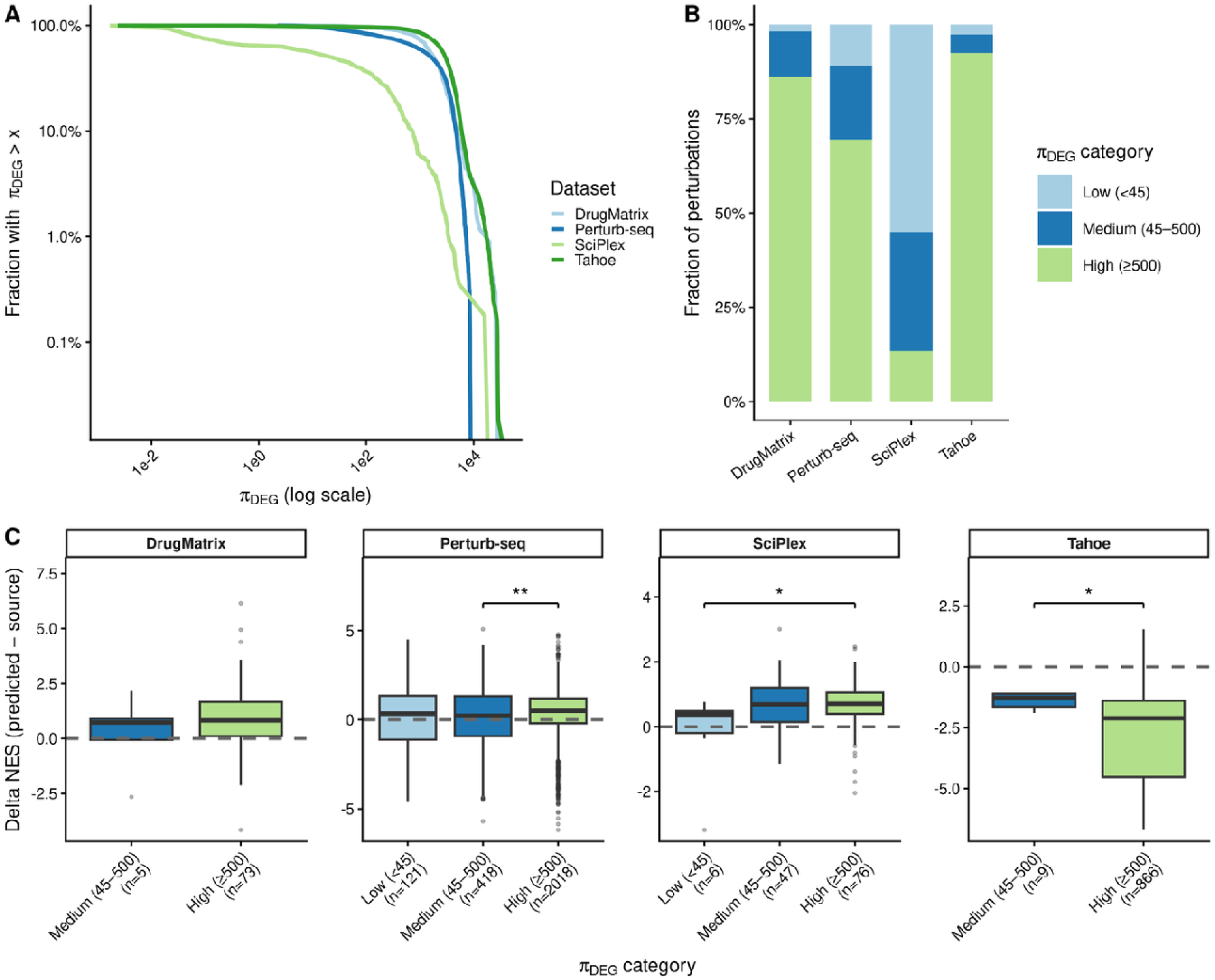
A) Complementary cumulative distribution plot of , both axes are in log scale. B) Fraction of perturbations in each category of . C) Delta NES (from the best performing method for each dataset) of perturbations grouped by categories for each dataset. Annotations between bars report statistical significance of a two-sided Wilcoxon test between groups e.g. p-value < 1e-2 indicated by **.

**Figure 6.**
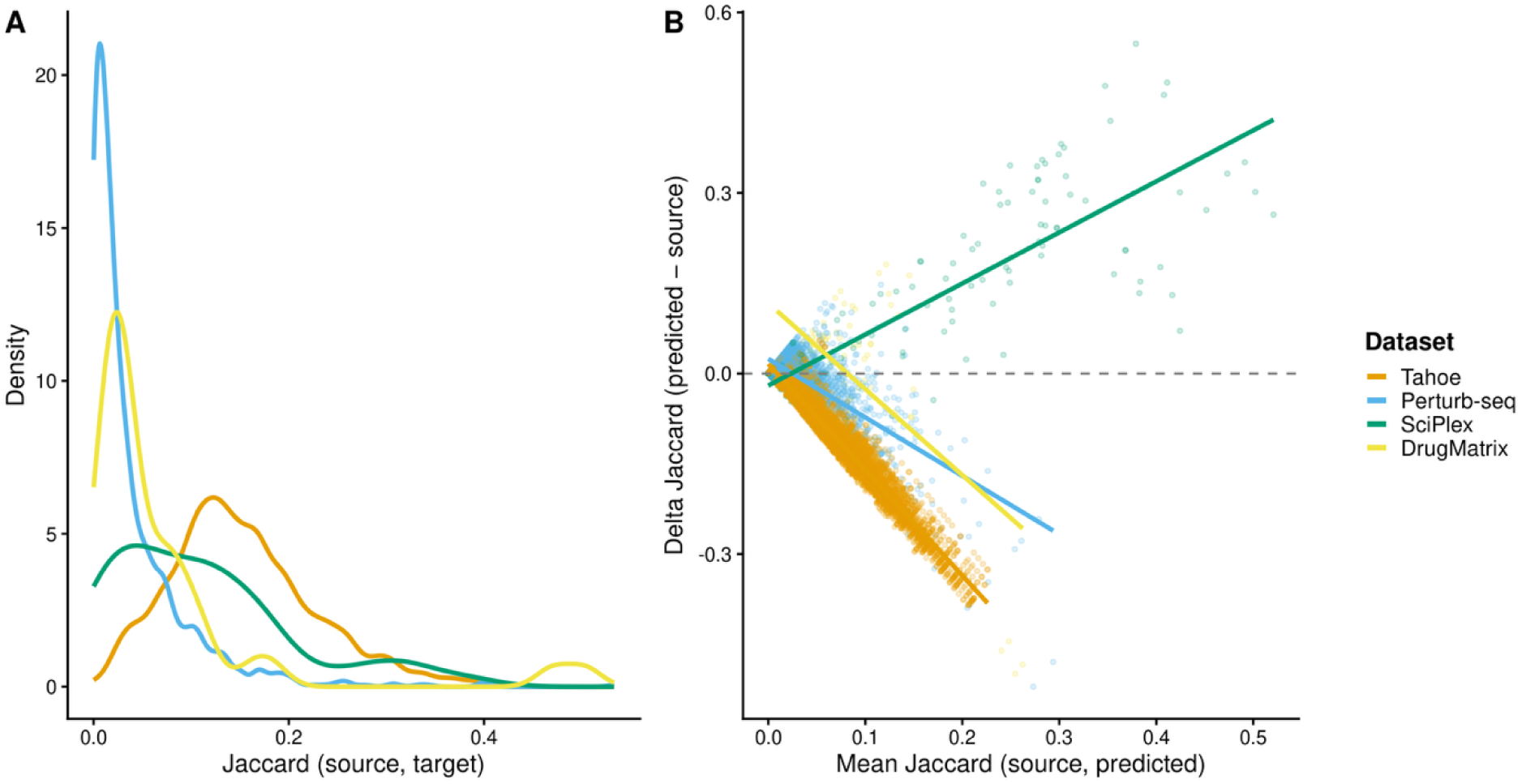
A) Distribution of Jaccard similarity for pairs of perturbational signatures for each dataset. B) Scatter of Delta Jaccard scores: [Jaccard (predicted, target) – Jaccard (source, target)] against Mean Jaccard similarity: Mean [Jaccard (predicted, target), Jaccard (source, target. Only the best method per dataset is reported: Tahoe, Perturb-seq – Net Prop, SciPlex – projCor (GSVA), DrugMatrix – Mean.

**Figure 7.**
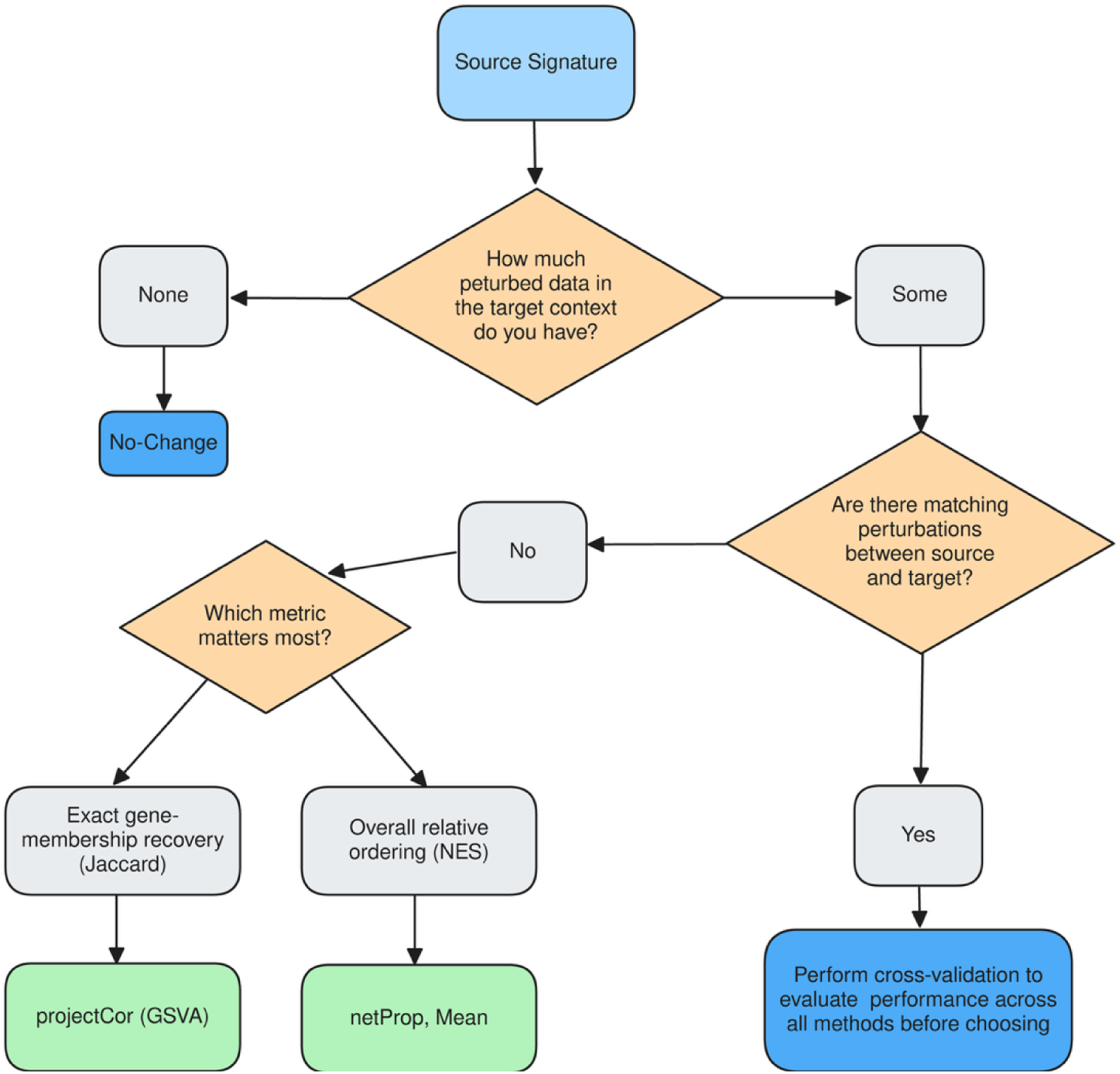
Practical decision workflow for selecting a recontextualization method. We recommend empirical validation against held-out target perturbations when available, since the best method is dataset and regime-dependent (Fig. 3).

#### Perturbational Strength

In addition, we investigated whether the overall statistical strength of perturbations is predictive of recontextualization performance. There are several statistical measures of the impact of perturbations, including, but not limited to, transcriptional activity score (TAS) of the L1000 Connectivity Map ^1^, energy distance ^4,37^, and mean squared error (MSE) between perturbed and control cells, but these are either defined on legacy assays such as L1000, or have ambiguous units in the case of E-Distance, or are optimized for model evaluation instead of perturbation evaluation in the case of MSE ^38^. We used TRADE, a recent method by Nadig et al., which estimates two complementary measures: 1) transcriptional impact (TI) – the variance of the LFC distribution; and 2) the effective number of differentially expressed genes (π*_DEG_*) which is defined as 3 *M*/*k* where M is the number of genes with measured expression, and *k* is the kurtosis of inferred LFCs) ^5^. These two statistical measurements are complementary as would distinguish between large TI driven either by a given perturbation having a large effect on a small subset of genes (high *K*, small π*_DEG_*), or if a perturbation results in small changes on a large subset of genes (small *K*, large π*_DEG_*).

We found that the distribution of the impact of perturbations varied across datasets and perturbations (Fig. 5a,b and Supp. Fig. 4). Except in SciPlex, most perturbations yielded a π*_DEG_*> 500 genes. Finally, we found that the correlation between perturbational impact and performance is dataset dependent, but largely positive. For instance, perturbations with a higher π*_DEG_* in Perturb-Seq and SciPlex were associated with better predictive performance (Fig. 5c), and perturbations with higher TI in SciPlex showed better predictive performance (Supp. Fig 4). It is important to note that SciPlex π*_DEG_* values did not converge and so should be interpreted with caution.

### Not all contexts are equal

Beyond perturbation-level properties, the difficulty of recontextualizing perturbation signatures varies substantially within datasets, depending on how similar the source and target biological contexts already are. We characterize this baseline similarity using the no-change metric distributions, specifically the Jaccard similarity and NES of the source perturbational signatures evaluated against the true target signatures, which quantify how well signatures transfer without any recontextualization. For instance, the Tahoe dataset exhibits a right-shifted distribution of both Jaccard and NES values relative to the other datasets, indicating that NCI-H23 signatures are already substantially similar to those measured in the target cell lines (Fig. 6a, Supp. Fig. 5a). To account for the coupling of the predicted and source similarity scores (both are compared to the target signature) we used Oldham’s method which compares the difference (Y-X) against the average (X+Y)/2 ^39^. Restricting data to the best-performing model per dataset (as identified in Fig. 3), we observe that the relationship between prediction performance (Delta Jaccard) and baseline context similarity (measured indirectly through Mean Jaccard) depends on the dataset. For SciPlex, there is a strong positive relationship between baseline similarity and prediction performance, i.e., similarity between the source signature and the target signature is correlated with the similarity between the *recontextualized* signature and the target signature (Fig. 6b). However, for *DrugMatrix*, Tahoe, and Perturb-seq baseline similarity is not correlated with recontextualized similarity, that is, as similarity between the source and target increases, similarity between the *recontextualized* signature and the target signature stays level or decreases (Fig. 6b). These dataset-specific trends suggest similarity between biological contexts (as measured by similarity of source and target perturbational signatures) is not always predictive of recontextualization performance. However, they do identify, for each dataset, which types of perturbations require more improvement, i.e., for *DrugMatrix* and *Perturb-Seq* perturbations that are not similar are being recontextualized well, but not those that are already similar, and for *Tahoe* all perturbations require improvement.

## Methods

### Perturbational signature derivation

For each of the perturbational datasets, we first derived differential expression signatures for each of the perturbations measured in each of the biological contexts (Table 1).

For the *Perturb-Seq* dataset, preprocessing was performed to remove low-quality cells according to three axes: total counts, number of unique genes, and mitochondrial gene expression. For the axes total counts and number of unique genes, we removed cells that were more than 5 median absolute deviations (MADs) from the median, and for mitochondrial gene expression, we removed cells that were more than 3 MADs from the median as recommended by best practices ^40^. Next, we generated pseudobulk count matrices using Scanpy’s Aggregate function ^41^ by summing counts across all cells belonging to each perturbation and gem_group within K562 and RPE1 separately. Differential expression was performed independently for each perturbed gene using DESeq2 ^42^, with a design of ∼ gem_group + perturbation_status, contrasting perturbed against non-targeting controls. Genes were ranked by the product of log fold change and negative log10 FDR-adjusted p-value, and the top 100 upregulated genes were retained per compound. Signatures were restricted to perturbations detected in both cell lines, and those with fewer than five significantly upregulated genes (FDR < 0.05, log2FC > 0) were excluded.

For the *SciPlex* dataset, single cell data was first preprocessed to remove outlier cells as defined by the axes of total counts, number of unique genes, and mitochondrial gene expression as above. Since there were multiple drug doses measured, we also filtered to the maximum tested dose for each compound. Pseudobulk profiles were then generated by summing counts within each product–replicate group using Seurat’s AggregateExpression function ^43^. Per-drug differential expression was computed with DESeq2 using a design of ∼ replicate + product_name, contrasting each drug against Vehicle controls. The same log fold change, p-value gene-ranking metric was then applied, and the top 100 upregulated genes were retained per compound. Signatures were restricted to the intersection of drugs detected across all three cell lines (A549, K562, MCF7), and perturbations with fewer than five upregulated genes were excluded.

For the *DrugMatrix* dataset, rat liver and kidney microarray ExpressionSets from the National Toxicology Program (NTP) were already preprocessed and log-transformed so no additional pre-processing was performed. Since multiple drug doses were present, we also subsetted each dataset to the maximum dose/drug with matching controls (same vehicle and time point). Using limma ^44^, a two-group linear model (Drug vs. Control) was fit with empirical Bayes moderation (eBayes), and FDR correction was applied via Benjamini–Hochberg. Genes were similarly ranked by the product of log fold change and −log adjusted p-value and the top 100 upregulated genes passing FDR < 0.05 were retained per compound. Signatures with fewer than five genes were excluded. Final signatures were restricted to drugs with qualifying results in both liver and kidney.

For the *Tahoe* dataset, we first preprocessed the single cell count data according to the same cutoffs mentioned in the original paper at least 700 unique molecular identifiers (UMIs), less than 20% mitochondrial reads, a UMI z-score within ±3 to remove outliers in total counts, a mitochondrial percentage z-score within ±3 to remove outliers in mitochondrial content, and at least 250 genes detected) ^2^. Then we generated pseudobulk AnnData objects by summing counts across cells belonging to the same sample, cell line, and drug treatment. The pseudobulk objects were likewise filtered to only include the highest tested dose per drug per cell line. For each drug–cell line combination, DESeq2 was run with design ∼ condition + plate, contrasting drug treatment against DMSO controls. Since the AnnData objects included a mix of HGNC symbols as well Ensembl IDs, identifiers were mapped to HGNC symbols using org.Hs.eg.db to maintain uniformity. The same log fold change, p-value gene-ranking metric was then applied, and the top 100 upregulated genes were retained per compound. Signatures were finally filtered to drugs present across all cell lines with at least five upregulated genes, and cell lines with fewer than five qualifying drugs were excluded from the benchmark. Since there are 50 cell lines in Tahoe and conducting benchmarking on all 50 choose 2 directions would have prohibitively high computational costs, we randomly sampled ten cell lines and all benchmarking was done from the source context NCI-H23 to the following nine target contexts: A498, HCT15, HEC-1-A, LoVo, MIA PaCa-2, Panc 03.27, SNU-1, SNU-423, and SW48.

### Baseline prediction methods

#### No-Change

The no-change baseline uses the source-context signature directly as the predicted target signature for each perturbation, performing no new prediction. This establishes the performance floor against which all methods are compared: any method must outperform it to demonstrate that cross-context adaptation adds value beyond simply transferring the source signature unchanged.

#### Mean

The mean baseline ignores perturbation-specific information entirely and instead predicts a single consensus signature from the target context, with the data used to build that consensus differing by regime, broadcast uniformly across all perturbations. In the control experimental regime, the mean signature was computed by obtaining a differential expression signature between two cell lines/tissues at baseline control states; in the low coverage (10% of perturbed target) and high coverage (90% of perturbed target) regimes, a mean signature was computed by contrasting perturbation vs. control in that subset of perturbed target cells. The mean baseline probes whether knowledge of the target context’s baseline transcriptional state, independent of any perturbation-specific signal, is sufficient to improve upon the no-change prediction.

### projectCor – gene set projection-based prediction

#### Conceptual framework

projectCor is based on the intuition that genes whose expression co-varies with a source signature’s activity in the target context are likely to participate in the same or related biological processes, even if the magnitude or identity of regulated genes differs between contexts. Rather than requiring explicit network learning or fine-tuning, projectCor leverages the inherent expression structure of the target dataset by identifying genes whose expression patterns best align with the signature’s projected activity (see Supp. Fig. 8). This approach assumes that context-specific regulatory wiring will be reflected in expression correlations, allowing adaptation of gene sets through statistical dependencies alone.

#### Technical approach

To estimate how perturbational signatures translate across biological contexts, we implemented a projection-based prediction method that derives recontextualized signatures from gene–signature activity correlations in a target dataset. Given a target expression matrix and a collection of input gene signatures, we first computed sample-level projection scores representing the activity of each signature in the target data. In all benchmarking regimes, the target expression matrix used for projection excluded the perturbational profiles that were being predicted. These projection scores can be derived using gene set variation analysis (GSVA) ^45^ or eigengene projection ^46^ based on the first principal component of standardized expression of the genes in the signature.

For each signature *s*, the projection score vector across samples is denoted proj_s_. For every gene *g* in the target dataset, we computed the Pearson correlation between its expression profile and the signature activity scores:

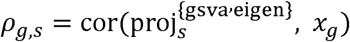

Where *x_g_* denotes the expression vector of gene *g* across samples. Genes were then ranked according to *p_g, s_*, and the top-ranked genes were selected to form a new predicted signature of the same size as the original gene set. Intuitively, this procedure identifies genes whose expression most strongly covaries with the projected activity of the source signature within the target biological context.

#### Implementation details

No hyperparameters required specification beyond the signature size, which was determined by matching the source signature. Predicted signatures were derived independently for each source-target pair and experimental regime. Compared to other methods benchmarked here, this method is computationally efficient, with typical runtime under one minute per dataset even for large gene expression matrices (>10,000 genes).

### netProp – network propagation-based prediction

#### Conceptual framework

netProp complements projection-based methods by leveraging gene co-expression network topology to extend predictions beyond genes with direct expression correlations to the source signature. The key insight is that genes functionally related to a perturbation often form connected communities in co-expression networks, even when direct correlation with the source signature is weak. By performing a guided random walk across a co-expression network learned from the target context, netProp discovers genes that are network-proximal to the source signature’s seed genes, capturing functional neighborhoods that may be obscured in pairwise correlation analysis (see Supp. Fig. 8). This enables adaptation of signatures through both topology and statistical significance.

#### Technical approach

We used weighted gene co-expression analysis (WGCNA) to learn gene-gene networks on the top variable genes of the target dataset (up to 10,000 genes) ^47^. In all benchmarking regimes, the target expression matrix used for network learning excluded the perturbational profiles that were being predicted. The resulting adjacency matrix is converted to a graph representation and normalized to produce a transition matrix governing diffusion across the network.

Given a binary seed matrix *P_0_* encoding the genes belonging to each input signature, we performed a random walk with restart to compute stationary probabilities that reflect the proximity of each gene to the seed set. At iteration *t*, the probability matrix *P_t_* was updated according to

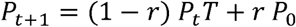

Where *T* is the normalized transition matrix and *r* is the restart probability. In implementation form, this update was computed as

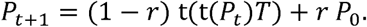

The algorithm iterates until convergence of the stationary probabilities. To reduce sensitivity to the choice of restart parameter *r* , we averaged results across a sequence of restart probabilities *r_1_*,…,*r_k_* . The final stationary score matrix was therefore

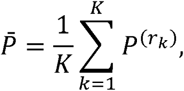

Where *P^(r_k_)^* denotes the stationary probabilities obtained using restart probability *r_k_*

#### Significance testing and gene selection

To identify genes significantly associated with each seed signature, we estimate empirical null distributions by performing bootstrap sampling of random gene sets matched by size to the original signatures. For each bootstrap replicat *b *∈* {1,…, B}* a random seed set was generated, and the same random-walk procedure was applied to obtain Gene-level significance was then estimated using the empirical percentile

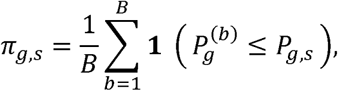

Where *P_g,s_ is* the stationary probability of gene *g* with respect to seed set *s*. Genes exceeding a specified percentile threshold were retained as members of the new predicted signature. In all experiments we used *B =* 30 bootstrap replicates. This procedure yields recontextualized signatures that reflect both the network topology and the statistical significance of gene proximity to the seed signature.

### Orthos

Orthos is an R/Bioconductor package that decomposes perturbational expression changes into context-independent (generic) and context-specific components using a pre-trained variational autoencoder. Orthos was trained on ARCHS4, and was designed to denoise biological context from perturbation signatures ^48^. As such, it serves as a context-independent baseline for evaluating whether other methods can successfully leverage context-specific information about a given perturbation. For each source context, the mean control expression profile (from DMSO or vehicle-treated cells) and a matrix of per-perturbation log2 fold changes were supplied to the *decomposeVar* function. Log fold changes were derived from DESeq2 for SciPlex and Tahoe, and from Limma for *DrugMatrix* (see Perturbational signature derivation). Orthos projects each perturbation contrast through its encoder and returns a residual contrast representing the context-independent signal. Genes were ranked by absolute residual magnitude, and the top N genes with positive residuals (matching the size of the corresponding source signature) were selected as the predicted target signature. We applied orthos to three datasets: *SciPlex*, *Tahoe*, and *DrugMatrix*. Orthos failed to produce signatures in the *Perturb-Seq* dataset due to issues resolving sufficient gene identifiers. Finally, since Orthos only decomposes perturbational signatures from differential expression tables and does not leverage additional data from the perturbed target, we only include this method in the benchmarking results for the control experimental regime.

### scGPT

scGPT is a single-cell foundation model pre-trained on 33 million normal human cells and can be fine-tuned for downstream tasks of batch-correction, cell type annotation, and perturbation prediction among other tasks ^7^. To adapt it for our benchmarking purposes, the model was initialized from the pretrained *scGPT_human* checkpoint, count matrices from *Perturb-Seq* were then subset to the gene features defined in scGPT, and lastly, we fine-tuned on 10 different splits of the perturbed target data from *Perturb-Seq* (K562 and RPE1) for both the low coverage and high coverage experimental regime. Unlike the other methods benchmarked, scGPT could not be evaluated in the Control regime as it needs to be fine-tuned on tuples of baseline-perturbed expression profiles. The gene universe for scGPT is limited to gene tokens encoded in the model and about 7,000 genes out of the 8,000 genes measured in the *Perturb-Seq* dataset could be predicted. Results reported in this analysis only apply to the low and high coverage regimes. This resulted in 40 scGPT models (ten splits of data for two target cell lines, across two types of benchmarking strategies). Each model was fine-tuned using default parameters (15 epochs, learning rate 1e-4, batch size 64). Since scGPT cannot predict chemical perturbations, without significant modifications, we only used it to benchmark on the *Perturb-Seq* dataset. Since, the scGPT output ends with a mean gene expression profile for each perturbation, we extended this method to derive a gene signature as output by first calculating z-scores for each gene against the target control distribution, applying FDR-adjustments per perturbation, and finally extracting the top ranked genes to obtain a recontextualized signature.

### STACK

STACK is a single-cell foundation model that uses in-context learning to generate predicted transcriptional profiles without task-specific fine-tuning ^12^. Given a *--base-adata* file of perturbed cells in a source context and a *--test-adata* file of unperturbed cells in a target context, STACK predicts counterfactual perturbed expression profiles in the target context. We applied STACK using the pretrained *bc_large_aligned* checkpoint via the stack-generation command-line interface, with default parameters (prompt-ratio 0.25, context-ratio 0.4, batch size 16, and 5 diffusion steps). For each of the datasets benchmarked (*Perturb-Seq, SciPlex, Tahoe*), source and target cell line AnnData objects were prepared and partitioned according to the three different benchmarking strategies (control, low coverage, and high coverage). The gene universe for STACK is also limited to the gene vocabulary encoded in the model (15,012 fixed panel) and for genes that could not be mapped, the model zero-pads the input expression profile. For each of the benchmarked datasets, STACK was able to predict 5,000-6,000 genes. STACK was pretrained on scBaseCount (approximately 149 million human cells from 10x Genomics runs mined from the Sequence Read Archive) and post-trained on a curated CELLxGENE subset together with the Parse PBMC cytokine dataset, and its authors used Tahoe-100M as a held-out zero-shot evaluation set ^12^. DrugMatrix is not part of this corpus. SciPlex (sci-RNA-seq3) is unlikely to be in scBaseCount, which comprises 10x Genomics data, whereas Replogle et al. Perturb-seq was generated with 10x Genomics and publicly deposited, so we could not verify whether those runs are present in scBaseCount. Methods also differ in their access to target-context data at inference: projectCor and netProp use the target-context expression matrix directly, STACK receives unperturbed target-context cells as prompts, and scGPT sees target-context data only during fine-tuning. Since STACK does not require fine-tuning, producing cross-validated predictions was just a matter of substituting different *--base-adata* files into the generation step. This mechanism of prediction differs from other foundation models like scGPT which require fine-tuning on new perturbational datasets. Further, since STACK was designed for single-cell data, and unlike the other datasets, *DrugMatrix* data was generated as bulk RNA sequencing data on a microarray platform, it was not included as a dataset for benchmarking. STACK’s output ends with an AnnData file containing predicted gene expression profile for each of the cells in *--test-adata.* We extended this method to derive a gene signature by calculating pseudo-bulk profiles and loading each into DESeq2 (contrasting predicted perturbed cells against matched control cells). The outputs of DESeq2 were then passed to the standard *sig_filter_fn* function to derive differential expression signatures.

### Evaluation

Each signature recontextualization method was evaluated by comparing its predicted signature against the true signature measured in the target context for the same perturbation. Two complementary metrics were used. Our first metric, Jaccard similarity, measures gene overlap between two signatures normalized by the size of their union. Our second metric was the normalized enrichment score (NES) from the *fgseaMultiLevel* function in the *fgsea* package. *fgsea* is a geneset enrichment scoring method which tests whether the predicted gene set is enriched at the top of the full differential gene expression target signature ^49^. NES captures how well the predicted genes rank among the most strongly upregulated genes in the target context, even when exact membership in the shortlisted signature differs. Both metrics were computed per perturbation and per cross-context mapping direction, yielding a paired distribution of predicted-versus-source scores for each method.

Using these paired scores, we assessed the statistical significance of improvement over the no-change baseline (only using the source signature as a prediction) using a one-sided paired Wilcoxon signed-rank test. Because each dataset typically involves multiple cross-context transfer directions (e.g., K562→RPE1 and RPE1→K562, or all pairwise cell-line pairs), a separate Wilcoxon test was performed for each direction. The resulting p-values were then combined across directions using Fisher’s method. Because transfer directions are not independent (reciprocal directions such as K562→RPE1 and RPE1→K562 use the same perturbations, and all Tahoe directions share NCI-H23 as the source context), the combined p-values were used only to rank methods and describe the consistency of improvement, not as formal statistical inference. The combined meta-p-value, as well as the median Wilcoxon test statistics are reported in Supp. Table 1. These paired scores summarize how well a method consistently improves upon the no-change baseline across all mapping directions within a dataset.

## Discussion

We benchmarked a suite of signature recontextualization methods across three experimental information regimes spanning four diverse perturbational datasets encompassing genetic and pharmacological perturbations in cell lines, as well as in vivo chemical perturbations in rat tissues. In the control regime, where only unperturbed target profiles were available, no method consistently outperformed the no-change baseline across any dataset, indicating that unperturbed expression alone is insufficient to predict the composition of perturbation-induced transcriptional changes. Results improved substantially once perturbed target profiles were introduced. In the low coverage and high coverage regimes, our projection-based and network propagation methods (*projCor, netProp*) emerged as consistently competitive, frequently matching or outperforming state-of-the-art foundation models including scGPT and STACK. Similar to previous benchmarking studies, we found that there is no ‘one-size-fits-all’ method, but that performance depends on target-data availability as well as perturbation type ^23^. However, our methods*, netProp and projCor,* are flexible across a variety of perturbational datasets: bulk or single-cell, chemical or genetic perturbations, unlike deep learning-based foundation models. In some contexts, the mean baseline, which estimates a single average perturbation signature across all perturbations, was still the strongest performer; this pattern was most pronounced in the DrugMatrix dataset, reinforcing the finding that simple baselines remain difficult to surpass in perturbation response prediction.

### Defining the recontextualization problem and establishing a rigorous evaluation framework

This work contributes methodologically by providing a precise definition of the *signature recontextualization* problem as well as a rigorous evaluation framework for benchmarking methods. Prior studies have lacked consistent problem formulations, and employed varying evaluation frameworks that precluded fair method comparison. We operationalize recontextualization as a sample-size-dependent learning problem: given a perturbation signature from a source context, we predict the set of differentially expressed genes when the same perturbation is applied to a target context, under varying constraints on reference data availability. Our evaluation paradigm instantiates this through three design choices: (1) systematically varying data availability in the target (control, low, high coverage regimes); (2) spanning diverse data types (genetic and chemical, single-cell and bulk, cell lines and tissues) to ensure results reflect genuine advances rather than modality-specific overfitting; and (3) applying consistent paired metrics (Jaccard similarity and NES) with rigorous hypothesis testing across all comparisons. This framework focuses method evaluation on the most pertinent outputs of perturbation modeling (predicting DEGs) and provides a scalable standard for positioning future methods quantitatively against a common benchmark.

### Uncovering the heterogeneity in perturbation predictability

Beyond method-level comparisons, our analyses also uncovered substantial heterogeneity in predictability in terms of properties pertaining to the perturbation itself, as well as the biological contexts that are involved. One key observation in the *DrugMatrix* dataset, was that statins and fibrates, which act through lipid-metabolism pathways broadly conserved between liver and kidney, tended to show more consistent recontextualization than corticosteroids and anthracyclines, whose mechanisms depend on tissue-specific regulatory or chromatin context (Fig. 4). One limitation of these results is that these classes comprise only 2–4 drugs each (statins 3, fibrates 4, corticosteroids 4, anthracyclines 2), these differences should be regarded as suggestive rather than definitive. We additionally observed MOA-level differences in *Tahoe* and *SciPlex*, but similarly these analyses were limited by sample size (Supp. Fig. 6, Supp. Fig. 7). A further limitation of the Tahoe analysis is that all nine transfer directions share NCI-H23 as the source context, so Tahoe effectively contributes a single source context; observations such as the lack of improvement over the no-change baseline in Tahoe may therefore reflect NCI-H23 specifically rather than Tahoe in general, and adding further source cell lines would address this.

Analysis of perturbational impact using the *π_DEG_ metric* confirmed that the distribution of perturbation strength varies across datasets: some datasets such as *SciPlex* mostly contain perturbations which affect only a small number of genes, while other datasets such as *Perturb-Seq, DrugMatrix*, and Tahoe mostly contain perturbations which drive broad transcriptome-wide changes. In *Perturb-Seq* and *SciPlex*, perturbations with a greater effective number of DEGs were more predictably mapped compared to those with much smaller impact, suggesting that methods benefit from stronger signal in the source context.

Another source of heterogeneity in perturbation predictability was the distribution of baseline context similarity: how similar source and target perturbations were to begin with. In *SciPlex*, perturbation pairs whose source and target signatures were already highly concordant showed the most improvement after recontextualization, however in the other three datasets: *DrugMatrix, Perturb-Seq*, and *Tahoe* the opposite was observed. Additional ways of quantifying similarity between biological contexts, beyond just the similarity of perturbational signatures, would be needed to further understand which recontextualization settings and contexts are most amenable.

### Considerations in deriving perturbational signatures

Deriving differential gene expression signatures requires a priori specifications of several parameters including but not limited to signature set size, DEG thresholds, and directionality, i.e., up or downregulated genes. For all the perturbational datasets, we applied uniform DEG thresholds, signature size cutoffs, and chose to only use upregulated genes (see Perturbational Signature Derivation section for details), and though these parameters were kept consistent for each dataset, it is important to note how these choices can affect downstream evaluation. For instance, signature size interacts differently with our two metrics: NES is normalized for gene-set size within fgsea, whereas Jaccard similarity is not, making Jaccard comparatively more sensitive to prediction errors in smaller signatures. Similarly, our focus on upregulated genes, biases our evaluation towards perturbations that result in gene induction rather than gene repression. We did not re-run the benchmark under alternative signature sizes, thresholds, or directionality choices, but note that these represent sources of variation that interact with the methods and could be explored in future extensions of this framework.

### Practical recommendations for method selection

Given that no single method consistently outperformed across datasets, benchmarking regimes, and evaluation metrics (Fig. 3), we summarize practical guidance for applying signature recontextualization to a new dataset as a decision workflow (Fig. 7). Users should first assess data compatibility: scGPT is restricted to genetic perturbations profiled in single-cell data, STACK requires single-cell input; by contrast, Mean, projectCor, and netProp are broadly applicable regardless of perturbation type or data modality. Users should then consider the availability of target data. In the control-only regime, none of the benchmarked methods reliably outperformed retaining the unmodified source signature (Fig. 2), so this remains the most defensible default in the absence of any target perturbation data. When some target data is available, Mean, projectCor-GSVA, and netProp are reasonable first candidates, as they were the methods that most consistently improved upon the no-change baseline across our benchmarked datasets (Fig. 3); however, the best-performing candidate differs by dataset and by evaluation metric. For instance, netProp ranked highest under NES for *Perturb-Seq*, while projectCor-GSVA ranked highest under Jaccard similarity for the same dataset — so users should weight candidates according to whether their application prioritizes overall ranking of genes (NES) or precise recovery of gene membership (Jaccard). Finally, when feasible, we recommend empirically validating candidate methods on available perturbations measured in both the source and target context, rather than relying on a fixed choice, since our results show that the identity of the best method is dataset- and regime-dependent.

Finally, the gene universe of each method is often overlooked, yet it is important to account for. For instance, many transformer-based foundation models are built on fixed gene vocabularies that may not include all genes measured in perturbational experiments. Other methods, such as STACK, may address this mismatch by zero-padding the input expression profile during inference, but this strategy can distort prediction performance. Thus, prediction models may need to be prioritized based on the size and fidelity of their internal gene vocabulary representations.

### Outstanding Challenges

A key challenge in perturbation prediction is accurately predicting the exact set of differentially expressed genes, as revealed by our signature recovery metrics: ranked enrichment and Jaccard similarity. Even for methods that show statistically significant improvement over the no-change baseline, gains in NES or Jaccard similarity remain modest at best. Some methods – notably projCor-GSVA and netProp – do achieve meaningful Jaccard gains in certain datasets; however, the overall magnitude of improvement remains low in absolute terms, highlighting that predicting the precise composition of perturbed gene sets across biological contexts remains an open problem.

Two properties of perturbational data which were highlighted in this study, compound this difficulty and illuminate where methods must improve most. First, for perturbations which result in a small number of differentially expressed genes, small errors in predicted gene membership have an outsized impact on Jaccard similarity and the signal available in a source signature is inherently limited. Second, current methods do not leverage baseline similarity between biological contexts in the prediction of perturbational signatures, and often struggle to recontextualize signatures in which the source and target signatures are already highly concordant – the cases for which prediction would seem most tractable. Third, none of the methods benchmarked here explicitly encode known biological priors. In our results we showed that recontextualization performance can depend on drug MOA. However, one could explicitly encode known drug target priors: whether a drug’s primary target is even expressed in a target cell line, or more generally, when recontextualizing perturbational signatures, whether a gene is even expressed in the target context. A drug whose primary target is absent from a new target context may be mechanistically unable to reproduce its source context effects no matter how well a recontextualization method captures transcriptional covariance, and source-signature genes that are simply silent in the target context cannot meaningfully inform a recontextualized prediction. Explicitly encoding these biological priors remains a promising direction for future methodological work.

Addressing these challenges will require advances on two fronts that go beyond the present benchmark. First, there is a pressing need to characterize the same perturbations not merely in pairs of biological contexts, but across many more, spanning diverse tissues, developmental stages, disease states, and species. As recently argued, virtual cell models need context, not just scale ^50,51^: without data that comprehensively samples the space of biological contexts, models will remain brittle to distribution shift, and the gains achievable by recontextualization will be fundamentally capped by the limited diversity of reference signatures. Second, perturbations do not act on the transcriptome in isolation and transcriptomic signatures alone may be insufficient to characterize and predict the full scope of a perturbation’s action. Perturbations that affect the proteome, or metabolome introduce regulatory changes that are partially or wholly undetectable at the mRNA level, yet information captured in one modality contains predictive signal for changes in others. Only by integrating measurements across multiple molecular layers will we be able to build the richer, cross-modal reference signatures needed to accurately map perturbational responses across contexts. Taken together, addressing the challenges of exact DEG prediction, improving sensitivity to sparse and context-similar perturbations, expanding the diversity of benchmarked biological contexts, incorporating multi-modal measurements, and leveraging existing biological annotations represent the most critical steps toward a truly generalizable virtual cell capable of recontextualizing signatures reliably across the full landscape of biological variation.

## Data Availability

Perturbational datasets were downloaded from the following sources:

- DrugMatrix, National Toxicology Program (NIEHS): https://ntp.niehs.nih.gov/data/drugmatrix
- Perturb-seq, processed Replogle et al. (2022) datasets via Figshare: https://plus.figshare.com/articles/dataset/_Mapping_information-rich_genotype-phenotype_landscapes_with_genome-scale_Perturb-seq_Replogle_et_al_2022_processed_Perturb-seq_datasets/20029387
- SciPlex, via Figshare: https://figshare.com/articles/dataset/sciPlex_dataset/24681285?file=43381398
- Tahoe-100M, raw data via Google Cloud Storage, as described by the Arc Institute: https://github.com/ArcInstitute/arc-virtual-cell-atlas/tree/main/tahoe-100M Pseudo-bulk expression matrices as well as perturbational signatures are available on Zenodo:
- DrugMatrix

`o` Pseudobulk: https://doi.org/10.5281/zenodo.21433031
`o` Signatures: https://doi.org/10.5281/zenodo.21432933
- Perturb-seq

`o` Pseudobulk: https://doi.org/10.5281/zenodo.21433138
`o` Signatures: https://doi.org/10.5281/zenodo.21432937
- SciPlex

`o` Pseudobulk: https://doi.org/10.5281/zenodo.21433011
`o` Signatures: https://doi.org/10.5281/zenodo.21432935
- Tahoe-100M

`o` https://doi.org/10.5281/zenodo.21433050
`o` https://doi.org/10.5281/zenodo.21433000

## Code Availability

Scripts for reproducing results and plots: https://github.com/montilab/sigRecon_Benchmarking Benchmarking R package: https://github.com/montilab/sigrecon

## Acknowledgements

We want to thank members of the longevity consortium for feedback regarding the design and evaluation of this framework, as well as Brian Cleary, Josh Campbell, Paola Sebastiani and Gerald Denis for insights and feedback throughout the development of this project.

## Author Contributions

A.C.: Conceptualization, Methodology, Software, Formal analysis, Investigation, Data curation, Writing – original draft.

S.M.: Conceptualization, Methodology, Supervision, Funding acquisition, Writing – review & editing. N.R: Writing – review & editing.

T.G.: Writing – review & editing.

N.S.: Writing – review & editing.

## Supplementary Figure Legends

**Supplementary Figure 1.**
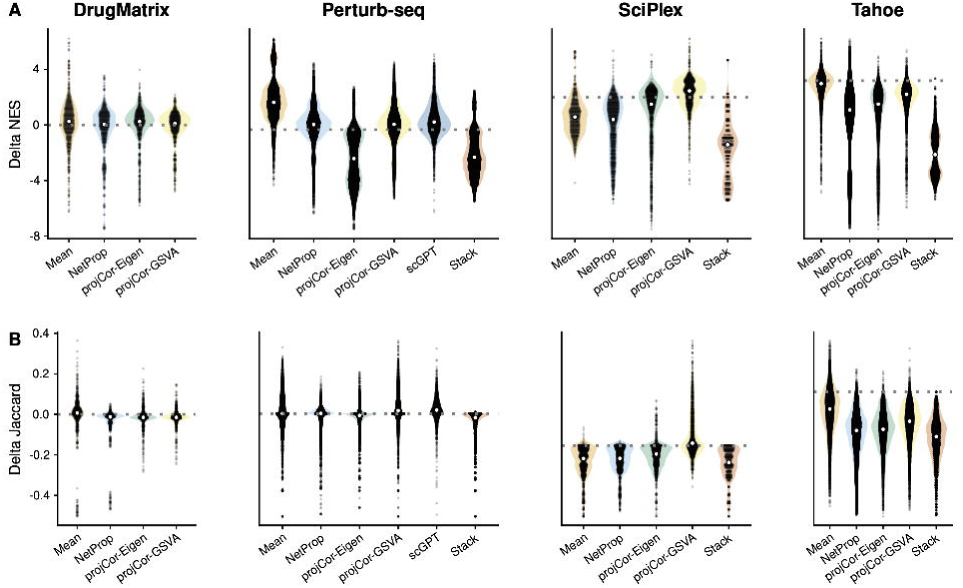
Performance of methods in the low coverage regime (1/10 of target perturbations available). A) Difference in NES between each method and the No-Change baseline [NES(predicted, target) − NES(source, target)] for each perturbation and transfer direction, by dataset. B) Difference in Jaccard similarity between each method and the No-Change baseline [Jaccard(predicted, target) − Jaccard(source, target)]. Values are medians across cross-validation splits of the perturbed target data. White points indicate the median of each distribution. scGPT was evaluated only on Perturb-seq, and STACK was not applied to DrugMatrix (see Methods).

**Supplementary Figure 2.**
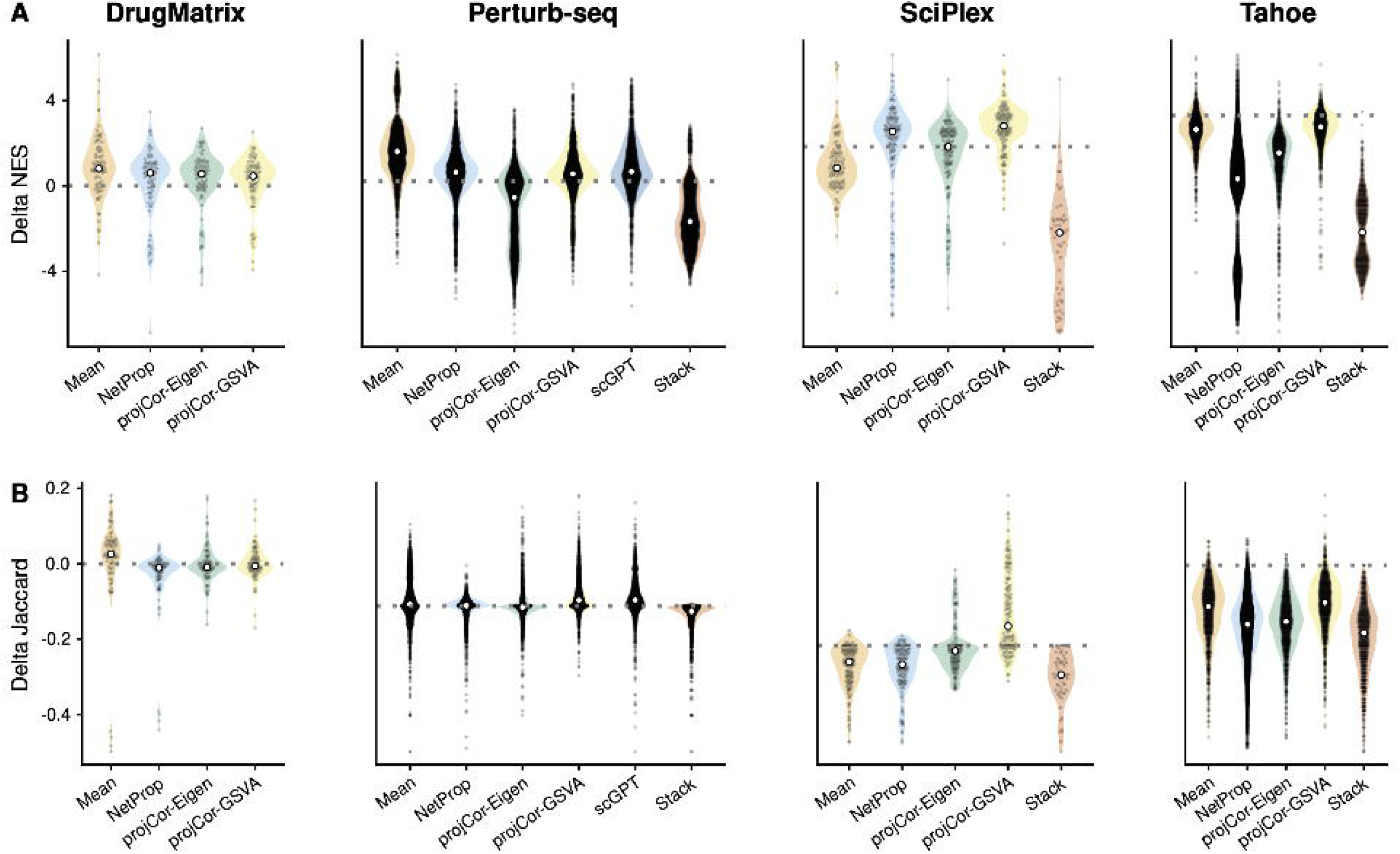
Performance of methods in the high coverage regime (9/10 of target perturbations available). A) Difference in NES between each method and the No-Change baseline [NES(predicted, target) − NES(source, target)] for each perturbation and transfer direction, by dataset. B) Difference in Jaccard similarity between each method and the No-Change baseline [Jaccard(predicted, target) − Jaccard(source, target)]. Values are medians across cross-validation splits of the perturbed target data. White points indicate the median of each distribution. scGPT was evaluated only on Perturb-seq, and STACK was not applied to DrugMatrix (see Methods).

**Supplementary Figure 3.**
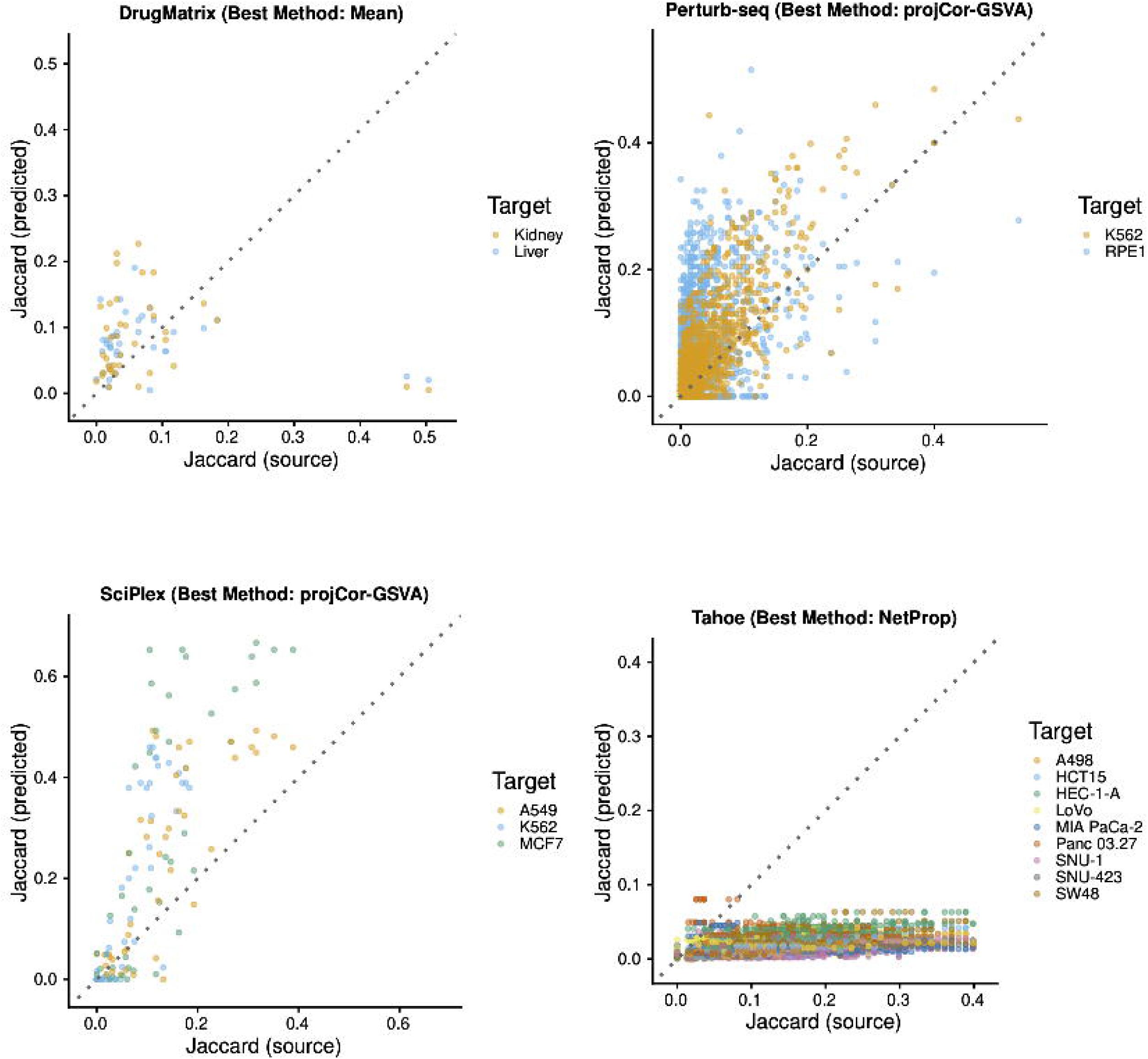
Jaccard similarity before and after recontextualization for the top-ranked method in each dataset. Each point is a perturbation in one transfer direction, colored by target context. The x-axis shows the Jaccard similarity between the source and target signatures (No-Change baseline) and the y-axis the Jaccard similarity between the predicted and target signatures. Points above the dotted identity line indicate improvement over the No-Change baseline. The top-ranked method by Jaccard similarity (Fig. 3A) is shown for each dataset: DrugMatrix, Mean; Perturb-seq and SciPlex, projCor-GSVA; Tahoe, netProp. Results are derived from the high coverage regime.

**Supplementary Figure 4.**
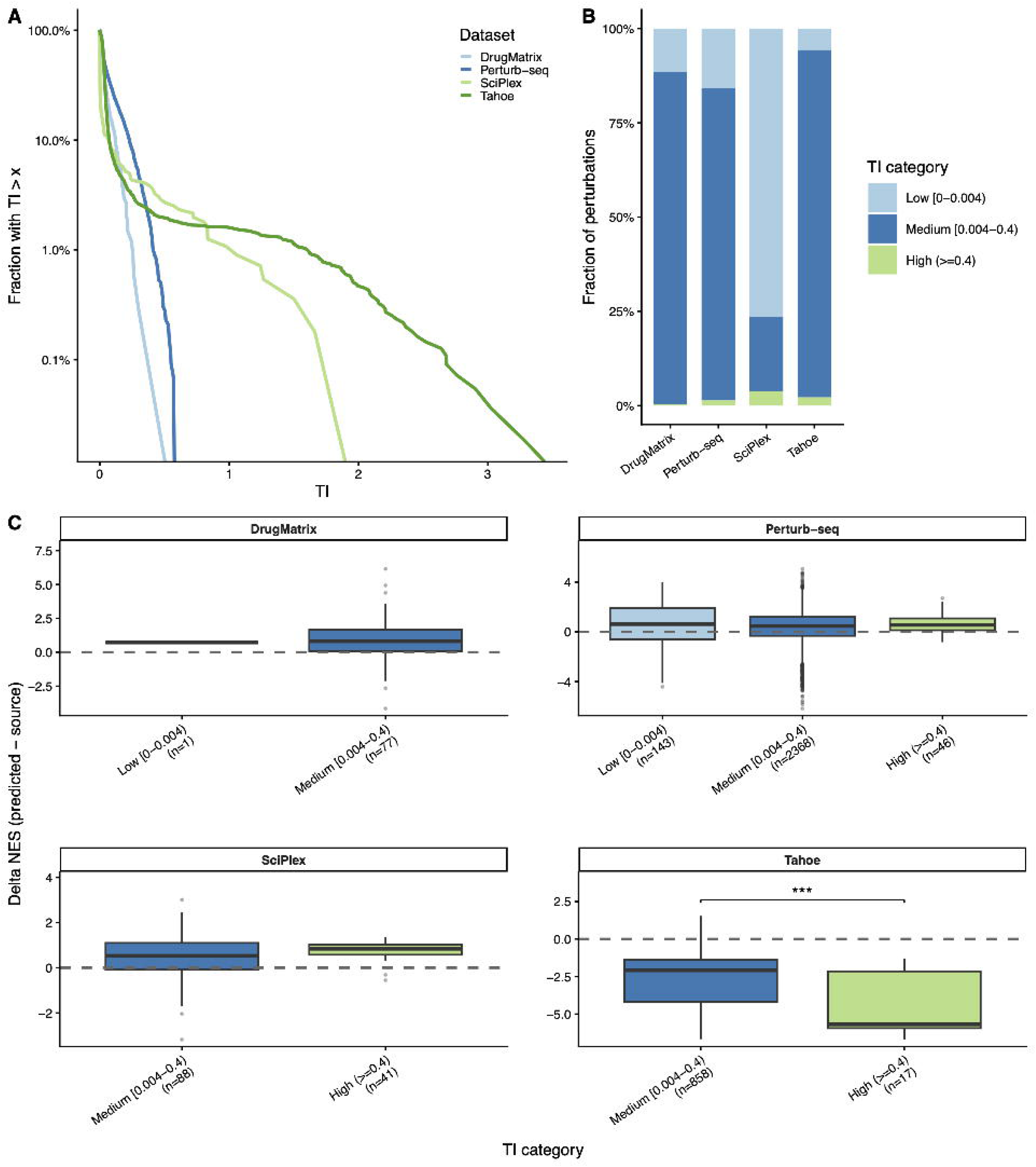
Transcriptional impact (TI) of perturbations and its relationship with recontextualization performance. A) Complementary cumulative distribution of TI (variance of the inferred log fold change distribution, estimated with TRADE) for each dataset; the y-axis is in log scale. B) Fraction of perturbations in each TI category: Low [0, 0.004), Medium [0.004, 0.4), and High (≥ 0.4). C) Delta NES [NES(predicted, target) − NES(source, target)] for the best-performing method in each dataset, grouped by TI category. Annotations between boxes report the significance of a two-sided Wilcoxon test between groups (*** p-value < 1e-3); comparisons without annotation were not significant.

**Supplementary Figure 5.**
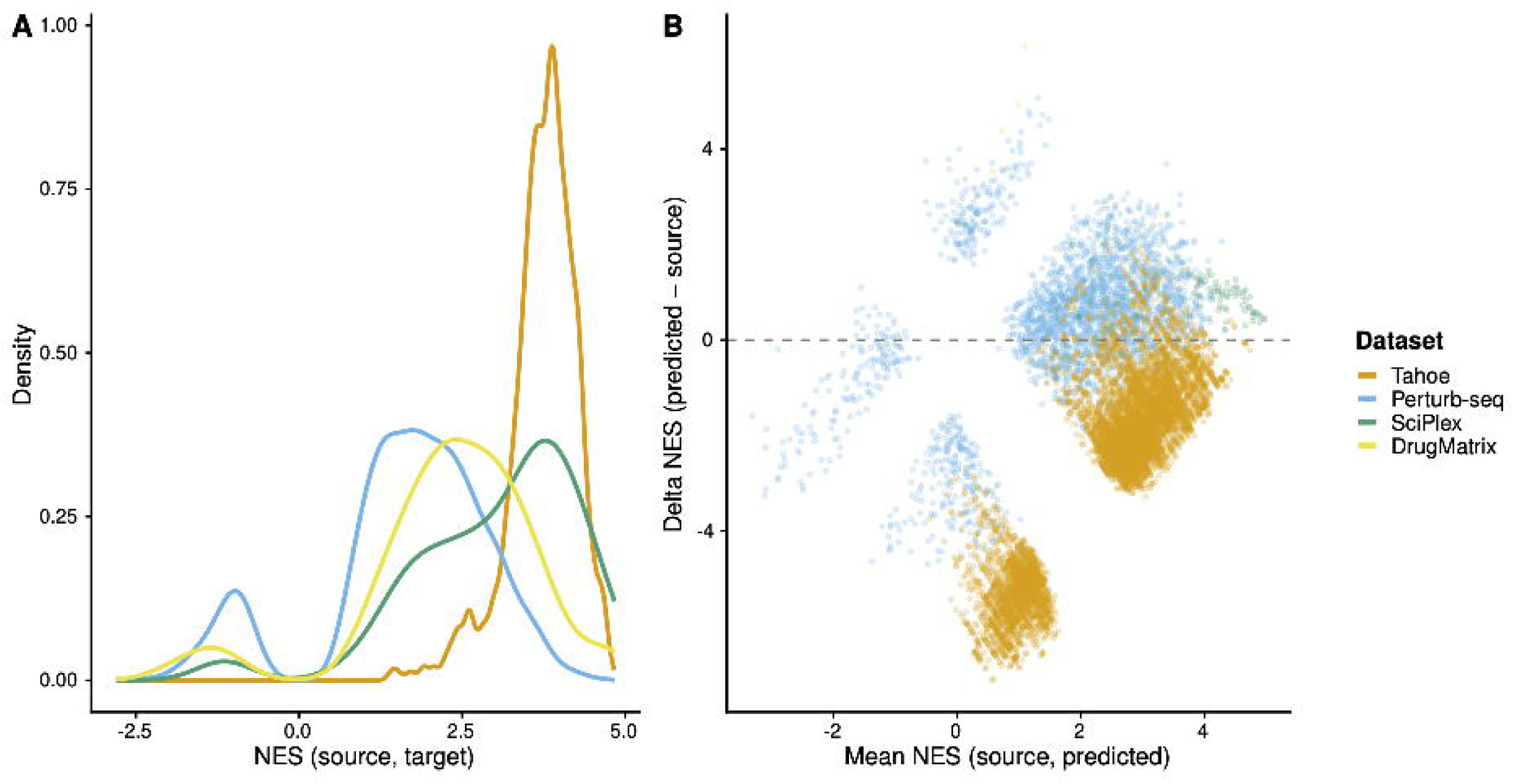
Baseline context similarity measured by NES and its relationship with recontextualization performance. A) Distribution of NES between source and target signatures (No-Change baseline) for each dataset. B) Delta NES [NES(predicted, target) − NES(source, target)] plotted against the mean of NES(predicted, target) and NES(source, target), following Oldham’s method to account for the coupling between the two scores. Only the best-performing method per dataset is shown: Tahoe and Perturb-seq, netProp; SciPlex, projCor-GSVA; DrugMatrix, Mean. The dashed line marks no change relative to the source signature.

**Supplementary Figure 6.**
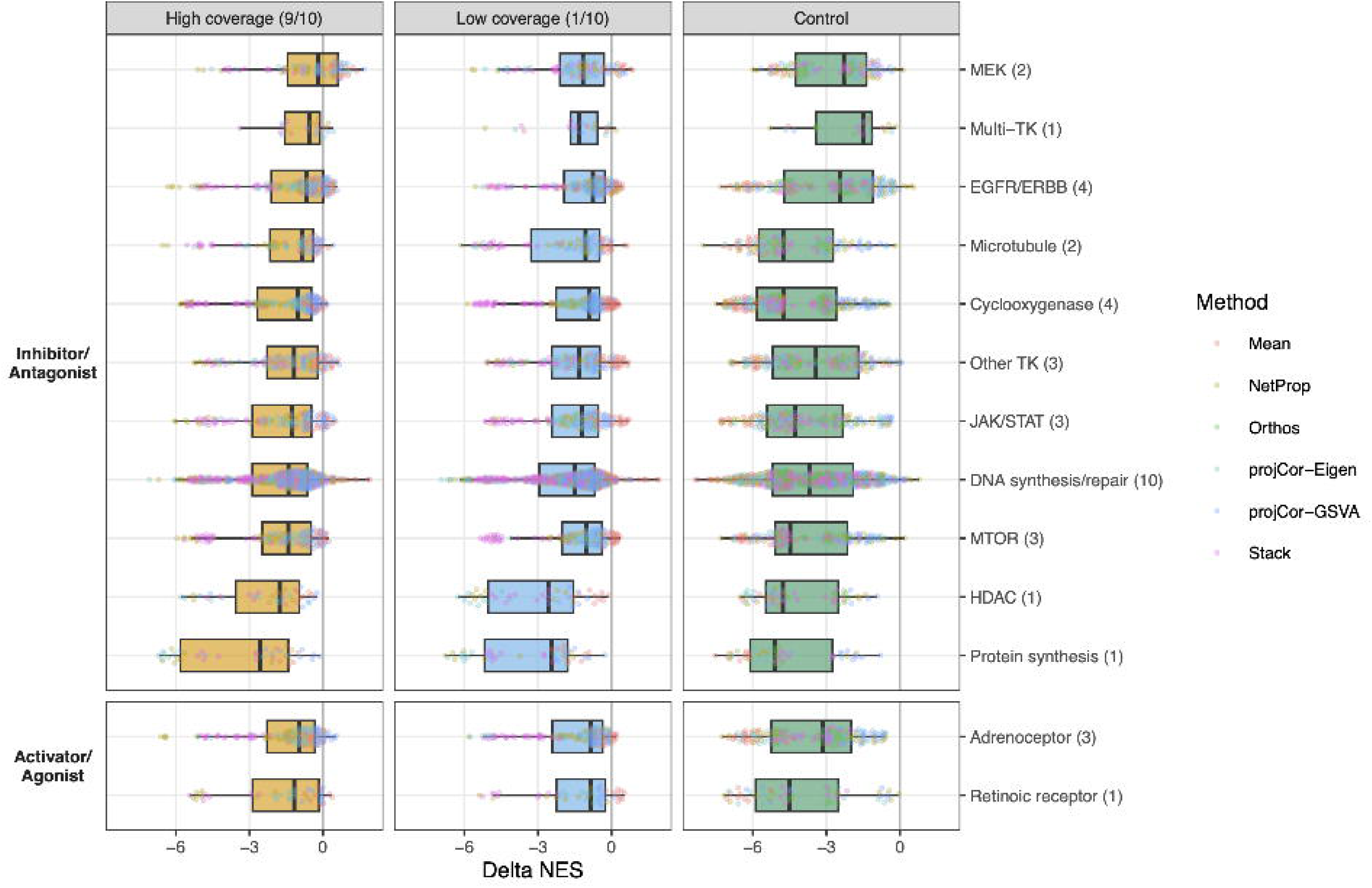
Recontextualization performance in Tahoe stratified by drug mechanism of action (MOA). Delta NES [NES(predicted, target) − NES(source, target)] for each drug, method, and target cell line, stratified by MOA class and benchmarking regime (high coverage, low coverage, control). MOA classes are grouped into inhibitors/antagonists and activators/agonists; numbers in parentheses indicate the number of drugs per class. Boxes show the median and interquartile range across all methods and target cell lines; points are colored by method. Most classes are represented by one to four drugs, so differences between classes should be interpreted with caution.

**Supplementary Figure 7.**
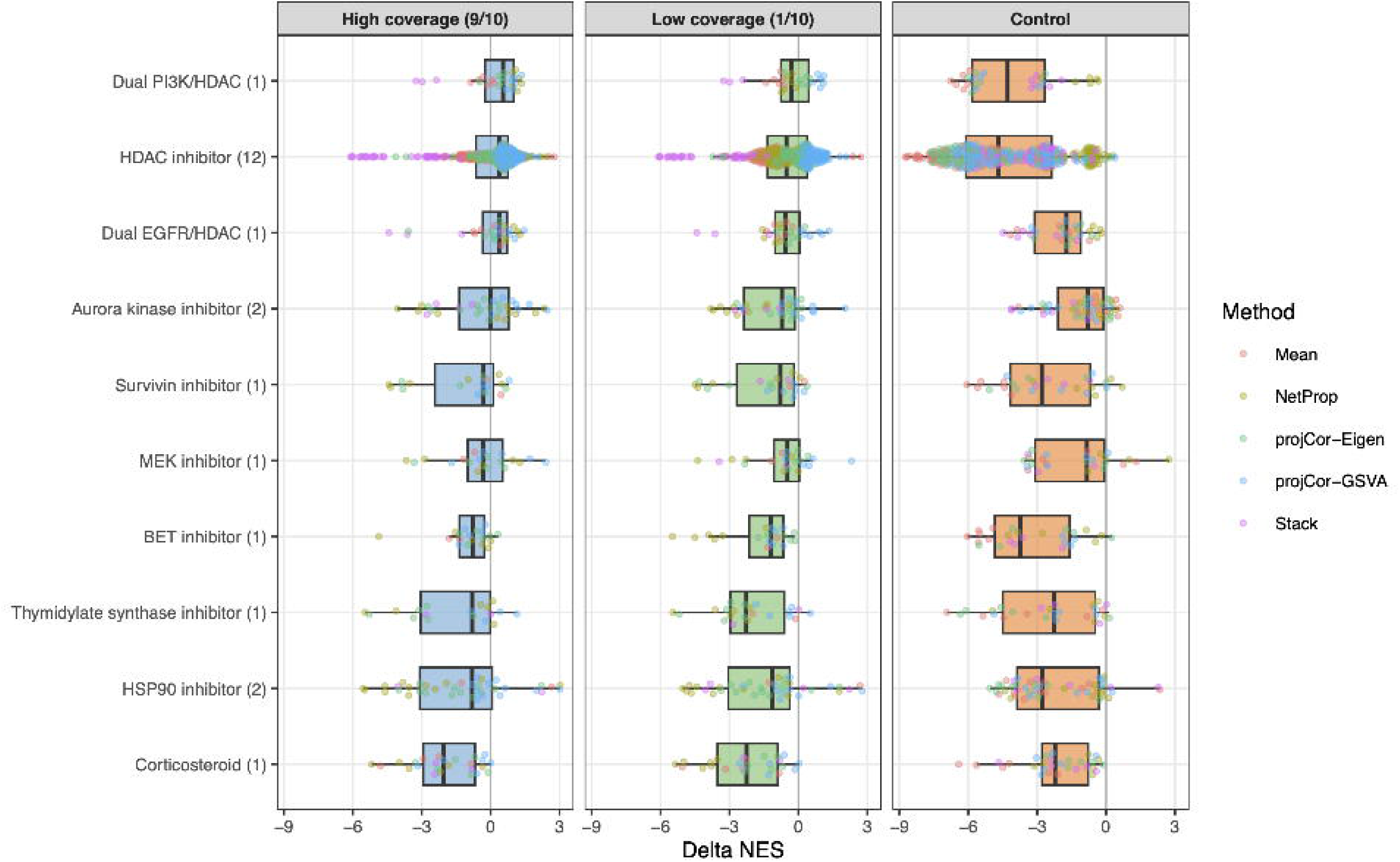
Recontextualization performance in SciPlex stratified by drug mechanism of action (MOA). Delta NES [NES(predicted, target) − NES(source, target)] for each drug, method, and target cell line, stratified by MOA class and benchmarking regime (high coverage, low coverage, control). Numbers in parentheses indicate the number of drugs per class. Boxes show the median and interquartile range across all methods and target cell lines; points are colored by method. Apart from HDAC inhibitors (12 drugs), classes are represented by one or two drugs, so differences between classes should be interpreted with caution.

**Supplementary Figure 8.**
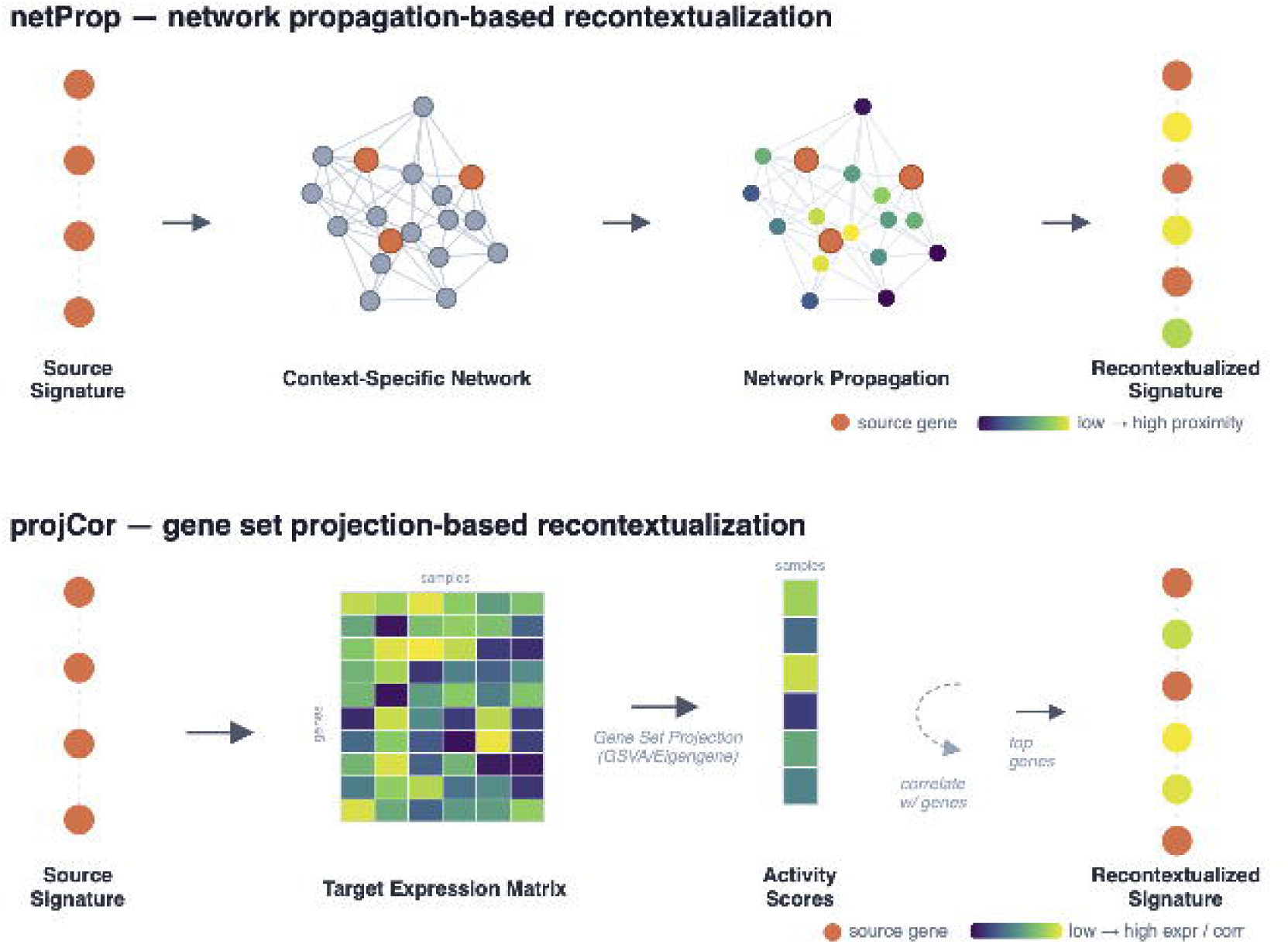
Schematic overview of the netProp and projectCor recontextualization methods. Top: netProp. Genes in the source signature (orange) are used as seeds for a random walk with restart on a co-expression network learned from the target context with WGCNA. Genes are scored by their network proximity to the seed set (color scale, low to high), assessed against size-matched random seed sets, and the top-scoring genes form the recontextualized signature. Bottom: projectCor. The source signature is projected onto the target expression matrix by GSVA or eigengene projection to obtain sample-level activity scores. Each gene’s expression is then correlated with these scores, and the top-ranked genes form the recontextualized signature, matched in size to the source signature.

